# Photosynthesis is led by stomatal circadian rhythm and not by starch depletion after a long period of darkness

**DOI:** 10.1101/2023.10.05.561115

**Authors:** Cédric Dresch, Véronique Vidal, Séverine Suchail, Huguette Sallanon, Florence Charles, Vincent Truffault

## Abstract

The circadian rhythm is an endogenous rhythm, defined by repeated metabolic oscillations every 24 hours under constant parameters. In plants, the circadian rhythm regulates the growth through photosynthesis and the management of carbohydrates. In indoor farming, the photoperiod is independent from the sunlight and is of interest to increase energy savings. In this work, we studied the effects of modifications of photoperiods on indoor-grown lettuces (*L. sativa*), which were then not tuned with the circadian rhythm. The pattern of photoperiods applied was: 16/12-16/8-16/8. A mutant-free study was carried out to avoid genes shutdown unintended side effects and to evaluate the impact of the circadian rhythm under marketable production conditions. We observed that the circadian regulation of the stomatal conductance was the main limiting factor of net photosynthesis when the photoperiod is exceptionally not tuned with the circadian rhythm. In our experiment, the disruption of the circadian rhythm decreased the photosynthetic activity by 6.2% throughout the light period, with no alteration of the yield, morphology or light and water use efficiencies. Furthermore, starch depletion induced by the changes in the photoperiod could not explain variations of the net photosynthetic activity during the following light period. Interestingly, disrupting the circadian rhythm saved 5% of lighting time compared to the control, which can be converted into energy savings. Consequently, the circadian rhythm can be defined as a limiting factor for reducing energy consumption in indoor farming. Further characterization of the links between the photosynthesis-carbohydrates-growth continuum and the circadian rhythm is thus required.

## Introduction

Photosynthesis is the main mechanism of carbohydrates production for plants and can be separated into two processes: light harvesting and CO_2_ fixation. Light harvesting process is defined as the absorption of sunlight photons and their conversion into chemical energy (Darwin, 1898; Samuelsson et al., 1983; Hennessey and Field, 1991). Light harvesting occurs in the photosystems, and mainly in the photosystem II (PSII). In fact, the light-harvesting complex of PSII is the most abundant integral membrane protein in chloroplasts and serves as the main solar energy collector (Liu et al., 2004). The absorbed photons initiate the photosynthesis, as they are necessary for water photolysis, NADP^+^ reduction and ATP biosynthesis (Caffarri et al., 2014). CO_2_ fixation occurs thanks to an influx of atmospheric CO_2_ into the leaf through stomata, regulated by stomatal conductance (Hennessey and Field, 1991). ATP, NADP+ and CO_2_ are then used to produce carbohydrates through the Calvin-Benson-Bassham cycle. The photosynthetic metabolism is influenced by environmental factors such as CO_2_ concentration, temperature and atmospheric humidity (Dodd et al., 2014b; Kaiser et al., 2015). As a matter of fact, they influence the most upstream limiting factors of photosynthetic activity, which are light harvesting efficiency, mainly photosystem II efficiency, and stomatal conductance. In addition to those well-known exogenous abiotic factors, more subtle endogenous factors are also involved in the regulation of photosynthesis, such as the circadian rhythm.

The circadian rhythm is an endogenous rhythm, defined by repeated metabolic oscillations every 24 hours under constant environmental parameters (Wang et al., 2022). The light harvesting process is mainly regulated by the circadian rhythmicity through the photosystem II. Indeed, post-translational regulation of the photosystem II occurs through reversible phosphorylation of D1 proteins, notably mediated by a circadian sensitive protein kinase (Bonardi et al., 2005). Moreover, 3 genes coding for the reaction center of photosystem II have been described as cycling (Harmer et al., 2000). Finally, the efficiency of photosystem II expresses daily variations that have already been measured on marine algae (*G. polyedra*), Crassulacean acid metabolism plants (*K. daigremontiana*), Brassicaceae (*A. thaliana*) and lettuces (*L. sativa)* (Samuelsson et al., 1983; Hennessey and Field, 1991; Rascher et al., 2001; Gould et al., 2009). It is relevant to note that as variations of the efficiency of the photosystem II were measured in marine organisms, they are specific to light-harvesting processes, *i.e.* not correlated to the stomatal conductance (Samuelsson et al., 1983). Stomatal conductance is directly linked to stomatal opening, which is known to be sensitive to light quality, as it is induced by blue and red light. Blue-light induction, defined as independent from the photosynthesis, has been precisely described to rely on a phosphorylation that activates H^+^-ATPases pumps, inducing an efflux of cytosolic H^+^ that leads to membrane hyperpolarization and K^+^ influx into the cell. This leads to an influx of K^+^ and Cl^-^, inducing a decrease in the hydric potential and H_2_O uptake in the vacuole of the cell guard. Consequently, cellular turgor pressure increases and leads to the opening of the stomata (Kinoshita and Shimazaki, 1999; Driesen et al., 2020). Unlike blue-light, red-light induction of the stomatal opening depends on the photosynthesis and may be linked to the electron transport rate (Messinger et al., 2006). Stomatal opening, whether it is induced by blue-light or red-light, depends on the light intensity (Zeiger, 2000; Driesen et al., 2020).

However, stomatal opening has also been described as specifically sensitive to the circadian rhythm. Indeed, free-running rhythmicity from 22 hours to 26 hours have already been described in *Vicia sp.*, *Arachis* and *Avena sp.* respectively (Pallas et al., 1974; Brogårdh and Johnsson, 1975; Gorton et al., 1989). The circadian regulation of stomatal opening is linked to specific circadian oscillators of the guard cells, already described in previous studies (Salomé et al., 2002; Hubbard and Webb, 2011; Hassidim et al., 2017). It can be noted that other parameters of photosynthetic activity, such as Rubisco activity and reactive Rubisco quantity, are not oscillating throughout the day in *P. vulgaris* and *L. polyedrum* and can be defined as insensitive to the circadian rhythm (Fredeen et al., 1991; Hollnagel et al., 2002). Put together, these data suggest that the daily variations of photosynthesis is mainly attributable to the circadian sensitivity of light harvesting processes and stomatal conductance. They therefore regulate the content of the direct end products of photosynthesis, carbohydrates.

Starch is one of the main end-products of photosynthesis (Sun et al., 1999; Stitt et al., 2010), as 30 to 50% of photosynthetic products are allocated into starch and other insoluble glucose polymers in *A. thaliana* (Stitt and Zeeman, 2012; Sulpice et al., 2014). Starch is an insoluble polymer accumulated linearly during the light period (Graf et al., 2010; Mengin et al., 2017; Flis et al., 2019), which plays a key role in the circadian sensitivity of carbohydrates (Sulpice et al., 2014). Indeed, the circadian clock regulates the rate of starch degradation so that 95% of the starch in *A. thaliana* is consumed during the night. This degradation rate can be calculated at the end of the day, anticipating the nighttime (Scialdone et al., 2013), in order to provide enough starch to last through the darkness period (Martins et al., 2013; Sulpice et al., 2014; Fernandez et al., 2017). The degraded starch produces sucrose (Aluko et al., 2021), a disaccharide defined as the main transportable form of carbon (Lemoine, 2000). Sucrose is known to affect circadian rhythms, notably through reactive oxygen species (Román et al., 2021), but to our knowledge, its specific relation with the circadian rhythm is less clear than that of starch. Consequently, starch is involved in the mobilization and management of carbohydrates and thus, in growth (Smith and Zeeman, 2020).

Growth depends on the circadian rhythm, which justifies to take it into account in agriculture and more specifically in indoor farming, where light signalization is independent from natural sunlight. In fact, when the photoperiod is not tuned with the circadian rhythm, productivity can be altered at least by 40% (Highkin and Hanson, 1954; Dodd, 2005). Moreover, it has been clearly described that maximum growth rates are reached when the plants are in the circadian resonance state, which occur when the photoperiod and the circadian rhythm are tuned (Dodd, 2005; Graf et al., 2010; Dodd et al., 2014a; Wang and Wang, 2015). Therefore, the circadian rhythm is a key component of the yield of plants. However, direct characterization of the link between circadian rhythm and growth is lacking at the moment and is crucial to reduce light and energy consumptions in indoor farming productions.

This study highlights the effects of a disruption of the circadian rhythm of plants in indoor farming conditions. In this case, the photoperiod and the circadian rhythm are exceptionally not tuned. Experiments were conducted on lettuces grown under a standard pattern of photoperiods (16/8 repeated continuously) or under a pattern of photoperiods with exceptionally long periods of darkness (16/12 – 16/8 – 16/8 repeated continuously). A mutant-free study was carried out to avoid genes shutdown unintended side effects, and thus to evaluate the importance of circadian rhythm under marketable production conditions.

## Material and methods

### Plant material and growth

Butterhead lettuce (*Lactuca sativa* L. var. fairly, ‘Enza Zaden’, The Netherlands) seeds were sown in 144 (16×9) holes germination trays filled with potting soil (TBSP, ‘Florentaise’, France). Water was provided until saturation by sub irrigation on a 3-days basis with a germination solution composed of 3.2, 0.6, 0.9, 1.3, 0.8 and 0.1 mmoles.L^-1^ of nitrogen, phosphorus, potassium, calcium, magnesium and sulfur respectively. Electroconductivity (EC) of the irrigation solution was equal to 0.5 mS.cm^-1^ with a pH of 5.6. Light intensity was set at 130 µmol.m^2^s^-1^ with Red:Blue 3:1 and ≈17% of white light, provided by LED lamps (T10 LED Grow Tube Light, HW-GL-T10-1200-36W-3Y, China). The photoperiod was 16/8 (16 hours of light, 8 hours of darkness) with respective temperatures of 24/19°C and relative humidity of 65%.

After 14 days, plantlets were transferred to a Gigrow® rotary culture system from Futura Gaïa Technology (figure 1A, France). Plants were rotating at a speed of 50 minutes per revolution, which provides a centrifugal force of 1,33×10^-5^ Newton, considered as neglectable. Lettuces were transplanted in 5 holes stainless steel trays (72 cm x 15 cm x 4.5 cm; length, width, height) with a density of 30 plants.m^-2^ filled with potting soil (VER4, ‘Florentaise’, France). Cultivation time was set for 31 days. Irrigation solutions were made with ‘Nutrimix^TM^’ of Futura Gaïa Technology. Temperature and vapor pressure deficit (VPD) were set at 24°C and 1KPa during the day and 19°C and 0.54 KPa during the night. Water supply was managed to saturate the soil, in order to avoid water deficiency effects (approximately 2.5 L.plant^-1^ for 31 days). Fertilization solution was composed of 29.3, 1.8, 12.5, 10, 7.5 mmoles.L^-1^ of nitrogen, phosphorus, potassium, calcium and magnesium respectively with an electroconductivity (EC) of 4 mS.cm^-1^ and a pH of 6.5. Overall light intensity for both modalities was equal to 350 ± 28 µmoles of photons.m^-2^.s^-1^ provided by ceramic metal-halide lamps (630W double ended, ‘Lumatek Ltd’, Malta). Light specter with a 4200K color temperature was provided, with a Red:Green:Blue ratio of 1:0.8:0.4. Two photoperiod modalities were studied (figure 1B):

- Control (CT) under a constant 16/8 photoperiod (16 hours of light/8 hours of darkness)
- Exceptionally long Period of Darkness (EPD) modality under a 16/12 – 16/8 – 16/8 photoperiods pattern.

**Figure 1.**
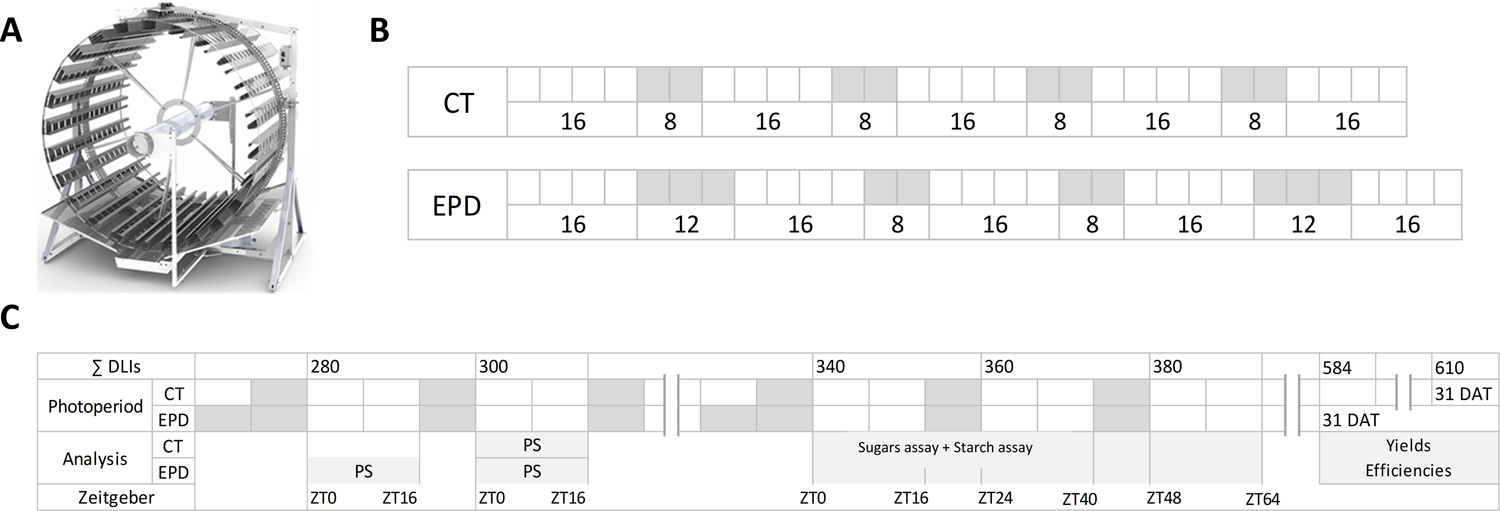
Cultivation system and light management during the experiment. (A) shows the GiGrow® cultivation system used in the experiment. (B) shows the photoperiod management in control lettuces, ‘CT’ and in the ‘Exceptionally long Period of Darkness’ modality, ‘EPD’. (C) shows the analyses conducted, scheduled according to the sum of DLIs (Daily Light Integrals), Zeitgebers (ZTs) and DAT (Days After Transplant). Analysis were: photosynthesis measurements (PS), sugars and starch assays and yield and efficiencies at harvest.

The analysis of samples was carried out according to two indicators: the sum of DLIs and the ZTs. DLIs are defined as the Daily Light Integrals, which represent the moles of photons (in the PAR) received each day by plants (moles of photons.m^-2^.d^-1^). ZTs are defined as “Zeitgeber”, which, in this experiment, represent the hours after the beginning of the light period (ZT0 = beginning of the light period).

*Photosynthetic measurement parameters: A_net_, Gsw, ΦPSII, F_v_’/F_m_’, and qP* Photosynthetic parameters are defined as follows (Mao et al., 2019):

*A_net_:* CO_2_ net assimilation rate (µmoles CO_2_.m^-2^.s^-1^), i.e. net photosynthetic activity.

*Gsw:* Stomatal conductance (moles H_2_O.m^-2^.s^-1^).

*F_s_:* Steady-state chlorophyll fluorescence level.

*F_0_’:* Minimum chlorophyll fluorescence yield in the light-adapted state.

*F_m_’:* Maximum chlorophyll fluorescence yield in the light-adapted state.

*F_v_’:* Maximum variable chlorophyll fluorescence yield in the light-adapted state.

*ΦPSII:* Quantum yield of PSII electron transport, calculated as (F*_m_*’-F*_s_*)/F*_m_*’.

*F_v_’/F_m_’:* Efficiency of excitation energy captured by opened PSII reaction centers, calculated as (*F_m_’-F_0_’*)/*F_m_’*.

*qP:* Photochemical quenching of chlorophyll fluorescence, calculated as (F*_m_’*-F*_s_*)/(F*_m_’*-F*_0_’*).

*A_net_, Gsw, ΦPSII, F_v_’/F_m_’* and *qP* were measured using a portable photosynthetic analyzer (Head version 1.4.7, LI-COR^®^ 6800, LincoEPD, USA) equipped with a LI-COR^®^ chamber type 6800-01. Parameters were set at 400 µmoles of photons.m^-2^.s^-1^ of light (Red and Blue light, R:B ratio of 9:1) with 420 ppm CO_2_, ambient temperature and humidity. Photosynthesis was measured at 300 DLIs for both modalities (after 15 or 16 days of growth) and also at 280 DLIs for EPD lettuces, which represents the light period after the exceptionally long night (Fig 1C). Photosynthesis was measured continuously for 16 hours, from the beginning of the light period (ZT0) until the beginning of the darkness period (ZT16). Measurements were taken manually with *A_net_*and *Gsw* values stability as recordable measurement indicators (3 to 5 minutes). Measurements were carried out on 10 plants (2 leaves per plant with the same phenological stage). During the measurements, plants were removed from the GiGrow^®^ culture system and the potting soil of plant were rehydrated every hour to avoid water deficit.

### Sugars analysis

Glucose, fructose, sucrose and starch contents were assessed at the end of the light period and of the darkness period at 340, 360 and 380 DLIs. Assays at the end of the darkness period correspond to ZT0, ZT24 and ZT48, while assays at the end of the light period correspond to ZT16, ZT40 and ZT64 for both modalities (figure 1C). In EPD lettuces, ZT0 corresponds to the beginning of the light period after an exceptionally long period of darkness.

### Glucose, fructose and sucrose contents by HPAEC

Lettuces were frozen in liquid nitrogen, grounded using IKA A11 basic Analytical mill (IKA^®^, Germany) and then freeze-dried using a Cryotec Cosmos freeze-dryer (Cryotec^®^, France). Between 15 and 20 mg of dried lettuces were sampled in ultra-pure water in order to obtain a concentration of 10 mg of DW.mL^-1^. Samples were homogenized and centrifugated 4 minutes at 15 000g, 4°C (Merck 3-16KL, KGaA^®^, Germany). Supernatant was collected and filtered through PTFE membrane filter (0.2 μm, IC Millex^®^-LG, Merck KGaA^®^, Germany). Glucose, fructose and sucrose contents were assayed using HPAEC (Dionex^®^ ICS-3000, USA) by injecting 5μL of each sample. Separation was carried out using an IC Dionex CarboPac^TM^ PA1 analytical anion-exchange column (10μm, 4 x 250mm, Dionex^®^, USA). The mobile phase was a solution of H_2_O (eluent A) and 250 mM NaOH with 4 mM sodium acetate (eluent B) with A/B ratio of 35:65 (v/v). An isocratic elution mode at a constant flow rate of 0.7 ml.min^-1^ was performed. Sugars were monitored by Pulsed Amperometric Detection (PAD), with a detection peak at 254 nm. Curves of glucose, fructose and sucrose contents were obtained by injecting calibration standards with concentrations of 2.5, 5, 7.5, 10, 12.5, 15 and 20 nmoles. CHROMELEON software v.6.7. was used to analyze the concentration of sugars. The limit of quantification was calculated according to Rusch et al. (Rusch et al., 2013).

### Starch extraction and starch content assays

Starch extraction was performed on 10 mg of dried lettuce. Samples were washed off of any free glucose twice with 1 mL of 90% ethanol. They were then centrifugated 2 minutes at 10 000g, 4°C, and 1 mL of ultra-pure water was added. Samples were autoclaved for 60 minutes at 121°C (ALV1131, ‘Getinge’, France) and weighted to determine the water loss during the autoclaving. 500 µL of ultra-pure water were added and samples were homogenized. Starch content was determined using a starch fluorometric assay kit (MAK368, Sigma-Aldrich, USA). Starch was hydrolyzed, revealed and starch content was determined by fluorimetry (Synergy HT, BioTek instruments, USA).

### Agronomical parameters

Fresh weight, dry weight and specific leaf area were studied at harvest (day 31). Fresh weight was measured using a precision balance XT 620M (PRECISA^®^, France). Two leaves were dried in an oven at 80°C for at least 3 days to calculate the dry weight. Specific Leaf Area (‘SLA’) was calculated based on leaf area per plant determined by ImageJ after picturing leaves in a 2D plan next to a standard meter (Schneider et al., 2012), using the following formula:

*Specific Leaf Area (cm^2^.mg^-1^ of DW) = (Leaf area per plant/Leaf dry weight per plant)* x 10^-3^

Light use efficiency (LUE) and water use efficiency (WUE) were calculated as follows:

*LUE based on the fresh weight (LUE FW, expressed as g of FW. moles^-1^ of photons) = Fresh weight/∑DLIs*

*LUE based on the dry weight (LUE DW, expressed as mg of DW. moles^-1^ of photons) = Dry weight/∑DLIs*

*WUE based on the fresh weight (WUE FW, expressed as g of FW.L^-1^) = Fresh weight/∑Volume of irrigation*

*WUE based on the dry weight (WUE DW, expressed as g of DW.L^-1^) = Dry weight/∑Volume of irrigation*

### Statistical analyses

Statistical analyses were carried on sugars analysis at each sampling time and on agronomical parameters values at 31 DAT to compare modalities, using the non-parametric pairwise Wilcoxon-Mann-Whitney test (n=5, α=0.05) on RStudio version 4.2.2 (Posit team, 2023).

## Results

### Daily photosynthesis declines earlier after an exceptionally long period of darkness

Photosynthesis was measured after a darkness period of 8 hours for control (CT) lettuces grown under a constant 16/8 photoperiod. For EPD lettuces, photosynthesis was measured directly after an exceptionally long darkness period of 12 hours, corresponding to the 2^nd^ light period of the 16/12 – 16/8 – 16/8 pattern (figure 1).

For control (CT) lettuces, *A_net_* values increased from ZT0 to ZT4 to reach a maximum of 10.6 ± 1.3 µmoles CO_2_.m^-2^.s^-1^. Then, *A_net_* values decreased to 9.0 ± 1.5 µmoles CO_2_.m^-2^.s^-1^ at ZT8 and to 4.4 ± 1.2 µmoles CO_2_.m^-2^.s^-1^ from ZT13 until the end of the light period (figure 2A). For EPD lettuces, *A_net_* reached a maximum of 10.4 ± 1.2 µmoles CO_2_.m^-2^.s^-1^ from ZT0 to ZT4 and decreased to reach a mean *A_net_* value of 5.5 ± 1.1 µmoles CO_2_.m^-2^.s^-1^ from ZT8 to ZT16 (figure 2B). When *A_net_* kinetics of CT lettuces and EPD lettuces were overlaid, we observed a 16% increase in *A_net_*values for EPD lettuces from ZT0 to ZT4 compared to the control (figure 2C). However, mean daily *A_net_* values were 6.2% lower for EPD lettuces compared to the control.

**Figure 2.**
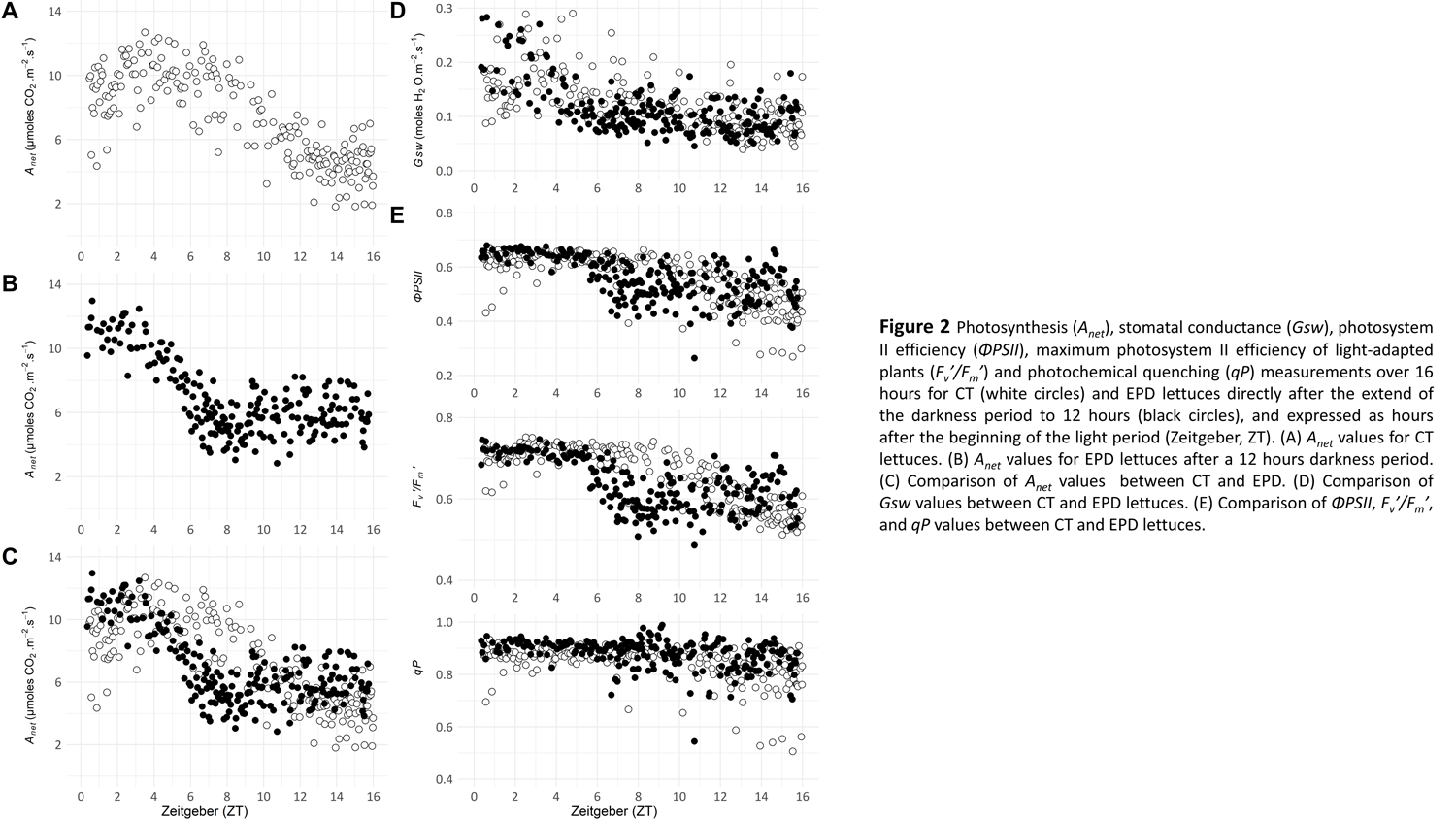
Photosynthesis (*A_net_*), stomatal conductance (*Gsw*), photosystem II efficiency (*ΦPSII*), maximum photosystem II efficiency of light-adapted plants (*F_v_’/F_m_’*) and photochemical quenching (*qP*) measurements over 16 hours for CT (white circles) and EPD lettuces directly after the extend of the darkness period to 12 hours (black circles), and expressed as hours after the beginning of the light period (Zeitgeber, ZT). (A) *A_net_* values for CT lettuces. (B) *A_net_* values for EPD lettuces after a 12 hours darkness period. (C) Comparison of *A_net_* values between CT and EPD. (D) Comparison of *Gsw* values between CT and EPD lettuces. (E) Comparison of *ΦPSII*, *F_v_’/F_m_’*, and *qP* values between CT and EPD lettuces.

*Gsw* values for CT lettuces increased from ZT0 to ZT2, peaking at a maximum mean value of 0.19 ± 0.07 moles H_2_O.m^-2^.s^-1^ between ZT3 and ZT4 (figure 2D). Subsequently, *Gsw* values decreased to reach a stable mean value of 0.09 ± 0.03 moles H_2_O.m^-2^.s^-1^ from ZT11 to ZT16. For EPD lettuces, *Gsw* maximum mean value of 0.20 ± 0.06 moles H_2_O.m^-2^.s^-1^ was measured from ZT0 to ZT3 and *Gsw* decreased to reach a stable mean of 0.10 ± 0.03 moles H_2_O.m^-2^.s^-1^ from ZT5 to ZT16. *Gsw* values increased in EPD lettuces of 7% from ZT0 to ZT4 compared to the control. However, mean daily *Gsw* values were 7.9% lower for EPD lettuces compared to the control.

For CT lettuces, *ΦPSII*, *F’_v_/F’_m_* and *qP* values were stable from ZT0 to ZT8 and equals to 0.62 ± 0.04, 0.71 ± 0.03, 0.87 ± 0.04, respectively (figure 2E). *ΦPSII*, *F’_v_/F’_m_* and *qP* values then decreased from ZT13 to reach stable means of 0.45 ± 0.07, 0.56 ± 0.03 and 0.79 ± 0.09, respectively. For EPD lettuces, *ΦPSII*, *F’_v_/F’_m_* and *qP* from ZT0 to ZT5 were stable and equals to 0.65 ± 0.02, 0.71 ± 0.01 and 0.91 ± 0.02 respectively. Then, *ΦPSII*, *F’_v_/F’_m_* and *qP* values decreased from ZT8 to reach stable means of 0.52 ± 0.06, 0.59 ± 0.04 and 0.87 ± 0.06 respectively. *ΦPSII* and *qP* values for EPD lettuces increased of 3% from ZT0 to ZT4 compared to the control. *F’_v_/F’_m_* values from ZT0 to ZT4 were similar in both modalities (less than 0.5% of difference). Mean daily *ΦPSII* values were similar in both modalities (less than 0.5% of difference), mean daily *F’_v_/F’_m_*values for EPD lettuces increased of 3.9% compared to the control and mean daily *qP* values for EPD lettuces decreased of 4.2% compared to the control. *A_net_* is closely linked to *Gsw*, *ΦPSII*, or *F’_v_/F’_m_*, and the duration of the darkness period did not influence those interrelations (Supplementary figure S1). However, *A_net_*, *Gsw*, *ΦPSII*, and *F’_v_/F’_m_*values decreased approximately 4 hours earlier for lettuces under EPD modality compared to the control.

### Daily photosynthesis declines in the same way as the control after returning to a standard period of darkness

We analyzed the photosynthesis after returning to a standard period of darkness. For the control, photosynthesis was still measured after a darkness period of 8 hours. For EPD lettuces, photosynthesis was measured after a standard darkness period of 8 hours following the exceptionally long period of darkness, corresponding to the third light period of the 16/12 – 16/8 – 16/8 pattern (figure 3).

**Figure 3.**
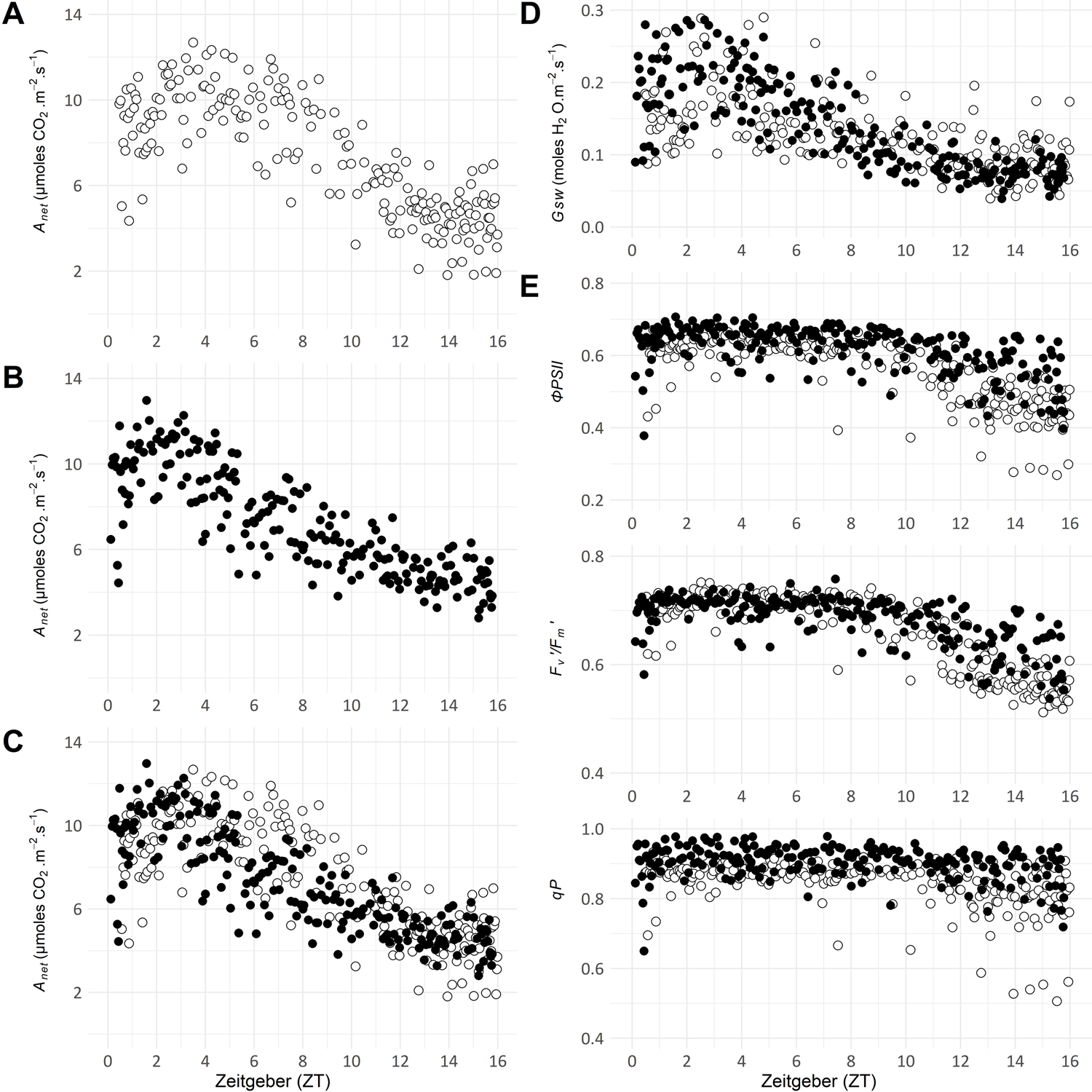
Photosynthesis (*A_net_*), stomatal conductance (*Gsw*), photosystem II efficiency (*ΦPSII*), maximum photosystem II efficiency of light-adapted plants (*F_v_’/F_m_’*) and photochemical quenching (*qP*) measurements over 16 hours for CT (white circles) and EPD lettuces after returning to a standard period of darkness (black circles), and expressed as hours after the beginning of the light period (Zeitgeber, ZT). (A) *A_net_*values for CT lettuces. (B) *A_net_* values for EPD lettuces after an 8 hours darkness period. (C) Comparison of *A_net_* values between CT and EPD lettuces. (D) Comparison of *Gsw* values between CT and EPD lettuces. (E) Comparison of *ΦPSII*, *F_v_’/F_m_’*, *qP* values between CT and EPD lettuces.

Control (CT) lettuces were grown under a constant 16/8 photoperiod. Lettuces experiencing an exceptionally long period of darkness (EPD) were grown under a pattern of photoperiods 16/12 – 16/8 – 16/8. Figure 3 shows photosynthesis-related data of CT lettuces after a darkness period of 8 hours (figure 3A) and of EPD lettuces after returning to a standard period of darkness of 8 hours (figure 3B).

*A_net_* values for EPD lettuces studied after a darkness period of 8 hours increased from ZT0 to ZT3 to reach a maximum value of 10.65 ± 0.94 µmoles CO_2_.m^-2^.s^-1^. *A_net_* values decreased to a stable mean value of 4.67 ± 0.81 µmoles CO_2_.m^-2^.s^-1^ from ZT13 to ZT16. When *A_net_* kinetics of CT lettuces and EPD lettuces were overlaid, the results were similar (figure 3C).

*Gsw* values for EPD lettuces increased from ZT0 to ZT3, peaking at a maximum mean value of 0.25 ± 0.06 moles H_2_O.m^-2^.s^-1^ and then decreased to reach a stable mean value of 0.08 ± 0.02 moles H_2_O.m^-2^.s^-1^ from ZT12 to ZT16 (figure 3D). When *Gsw* kinetics of CT lettuces and EPD lettuces were overlaid, the results were similar.

*ΦPSII*, *F’_v_/F’_m_* and *qP* values from ZT0 to ZT8 were stable and mean values equal to 0.65 ± 0.05, 0.70 ± 0.03 and 0.91 ± 0.04 respectively (figure 3E). *ΦPSII*, *F’_v_/F’_m_*and *qP* values then decreased until ZT13 to reach stable means of 0.55 ± 0.07, 0.62 ± 0.05 and 0.88 ± 0.06 respectively. As for *A_net_*and *Gsw*, *ΦPSII*, *F’_v_/F’_m_* and *qP* daily trends for EPD lettuces were similar to the control when EPD lettuces were exposed to a darkness period of 8 hours. To conclude, no differences in *A_net_*, *Gsw*, *ΦPSII*, *F’_v_/F’_m_* and *qP* daily patterns have been observed between CT lettuces and EPD lettuces after returning to a standard period of darkness.

### Glucose, fructose, sucrose and starch regulations respond to an exceptionally long period of darkness

Glucose content for CT lettuces was stable from ZT0 to ZT64 with a mean value of 100.2 ± 8.7 mg. g^-1^ of DW (figure 4A). No statistical differences have been found between ZT0 and ZT64. In EPD lettuces, glucose content values varied between 66.4 ± 5.8 mg. g^-1^ of DW at ZT48 and 95.6 ± 9.2 mg. g^-1^ of DW at ZT64. Therefore, the glucose content was significantly lower in EPD lettuces at ZT0, ZT16, ZT24, ZT40, and ZT48 compared to the control.

**Figure 4.**
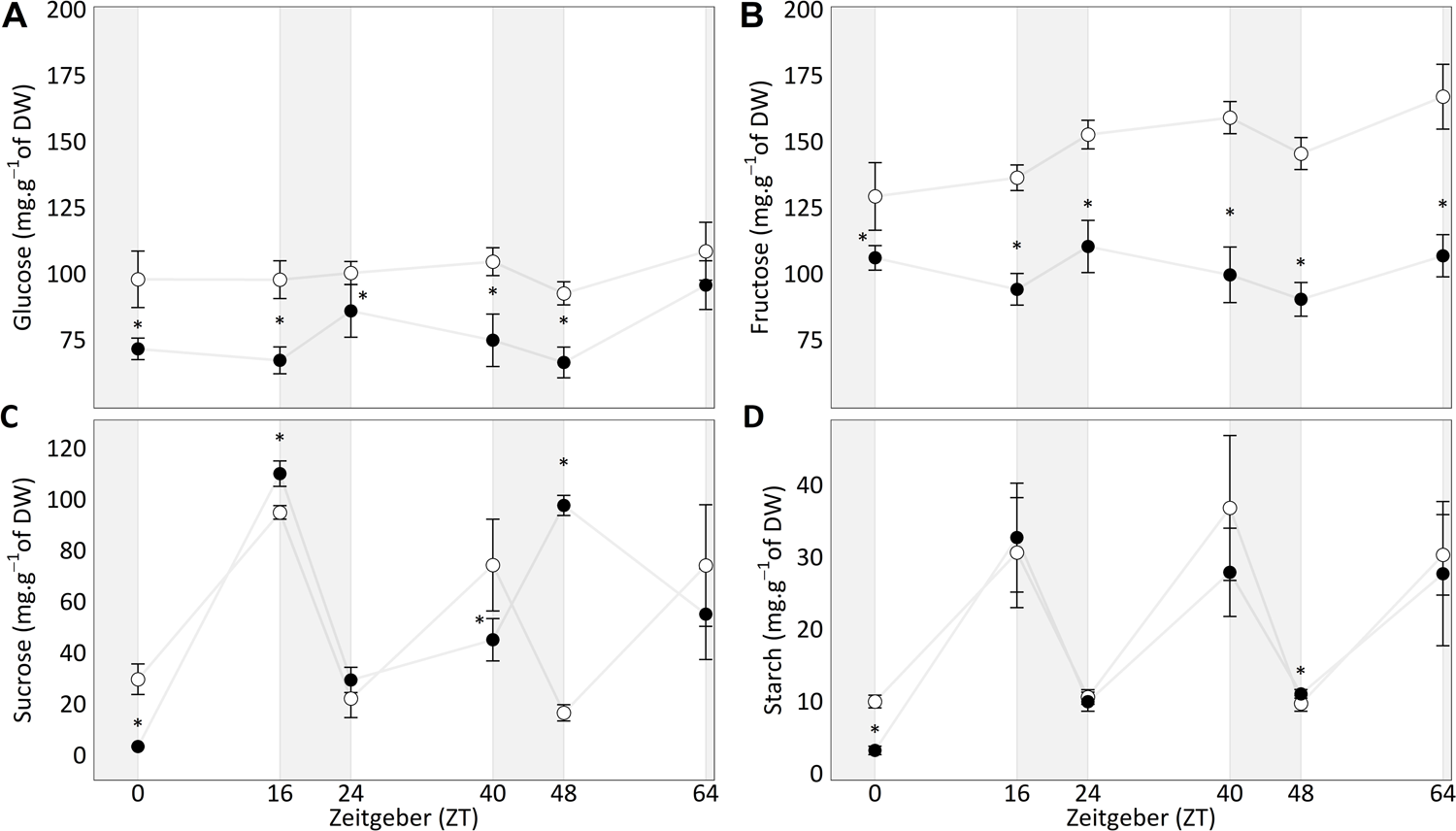
Carbohydrates contents over 3 consequent days on Control lettuces (CT) and EPD lettuces (Extended darkness period). For EPD lettuces, ZT0 represents a study after a period of darkness of 12 hours and for CT after 8 hours of darkness. (A) Glucose content. (B) Fructose content. (C) Sucrose content. (D) Starch content. Significant differences at the pairwise Wilcoxon-Mann-Whitney are represented with stars (n=5, α=0.05).

Fructose content for CT lettuces increased from 129.2 ± 12.8 mg. g^-1^ of DW at ZT0 to 166.9 ± 12.3 mg. g^-1^ of DW at ZT64 (figure 4B). Values for EPD lettuces were between 90.2 ± 6.4 mg. g^-1^ of DW at ZT48 and 110.2 ± 9.9 mg. g^-1^ of DW at ZT24. In EPD lettuces, no significant differences in fructose content have been found between ZT0 and ZT64. Unlike the control, the fructose content did not increase in EPD lettuces. Moreover, the fructose content was significantly lower than the control in EPD lettuces at all sampling times (ZT0, ZT16, ZT24, ZT40, ZT48 and ZT64).

The sucrose content in lettuces changes depending on the photoperiod (figure 4C). For CT lettuces, sucrose content values at the end of the darkness period were equal to 29.5 ± 6.0 mg. g^-1^ of DW, 22.0 ± 7.5 mg. g^-1^ of DW and 16.42 ± 3.2 mg. g^-1^ of DW at ZT0, ZT24 and ZT48 respectively. Sucrose content values at the end of the light period were equal to 94.7 ± 2.7 mg. g^-1^ of DW, 74.1 ± 18.0 mg. g^-1^ of DW and 74.0 ± 23.8 mg. g^-1^ of DW at ZT16, ZT40 and ZT64 respectively. For EPD lettuces, sucrose content values at the end of the darkness period were equal to 3.2 ± 0.3 mg. g^-1^ of DW, 29.3 ± 4.9 mg. g^-1^ of DW and 97.5 ± 4.0 mg. g^-1^ of DW at ZT0, ZT24 and ZT48 respectively. Sucrose content values at the end of the light period were equal to 109.9 ± 5.0 mg. g^-1^ of DW, 45.0 ± 8.3 mg. g^-1^ of DW and 55.0 ± 17.8 mg. g^-1^ of DW at ZT16, ZT40 and ZT64 respectively. For EPD lettuces, sucrose content values at ZT0 are 9.2-fold lower than the control. At ZT16 and ZT24, the sucrose content is equivalent to the control. From ZT40 to ZT48, the sucrose content for EPD lettuces increased to reach a 5.9-fold higher content than the control at the end of the darkness period. In the end, from ZT48 to ZT64, the sucrose content for EPD lettuces decreased to reach a similar value as the control at the end of the light period.

The starch content of lettuces changes depending on the photoperiod (figure 4D). For CT lettuces, starch content values at the end of the darkness period were equal to 9.92 ± 0.9 mg. g^-1^ of DW, 10.6 ± 1.0 mg. g^-1^ of DW and 9.7 ± 1.1 mg. g^-1^ of DW at ZT0, ZT24 and ZT48 respectively. Starch content values at the end of the light period were equal to 30.6 ± 7.6 mg. g^-1^ of DW, 36.8 ± 10.1 mg. g^-1^ of DW and 30.3 ± 5.6 mg. g^-1^ of DW at ZT16, ZT40 and ZT64 respectively. For EPD lettuces, starch content values at the end of the darkness period were equal to 3.2 ± 0.6 mg. g^-1^ of DW, 9.9 ± 1.3 mg. g^-1^ of DW and 10.1 ± 0.6 mg. g^-1^ of DW at ZT0, ZT24 and ZT48 respectively. Starch content values at the end of the light period were equal to 32.7 ± 7.5 mg. g^-1^ of DW, 27.8 ± 6.1 mg. g^-1^ of DW and 27.7 ± 10 mg. g^-1^ of DW at ZT16, ZT40 and ZT64 respectively. For EPD lettuces, starch content values at ZT0 are 3.2-fold lower than the control. At ZT16, ZT24, ZT40 and ZT64, starch values for EPD lettuces were similar to the control. In this experiment, the starch content increased during the light period and decreased during the darkness period in both modalities. Moreover, starch is more degraded under a longer period of darkness (12 hours) than under a standard period of darkness (8 hours).

### Agronomical traits at harvest are not affected by exceptionally long periods of darkness during growth

Fresh weight (FW), dry weight (DW), Specific Leaf Area (SLA), Light Use Efficiencies (LUEs) and Water Use Efficiencies (WUEs) have been studied at harvest (after 31 days) (Table 1). No statistical difference of yield per plant or efficiencies for a single plant have been found between modalities. However, a decrease of light of approximately 5% was measured in EPD lettuces compared to the control. We also calculated the LUEs for the production of an entire cultivation system (Jin et al., 2022). This LUE represents the fresh or dry matter divided by the surface of cultivation and the sum of DLIs. For CT lettuces, it was equal to 13.3 g of FW.m^-2^.∑DLIs^-1^ and 0.4 g of DW.m^-2^.∑DLIs^-1^. LUEs for EPD lettuces were equal to 15.2 g of FW.m^-2^.∑DLIs^-1^ and 0.5 g of DW.m^-2^.∑DLIs^-1^. With our experimental set-up, we could not calculate the standard deviations of these light-use efficiencies.

**Table 1.**
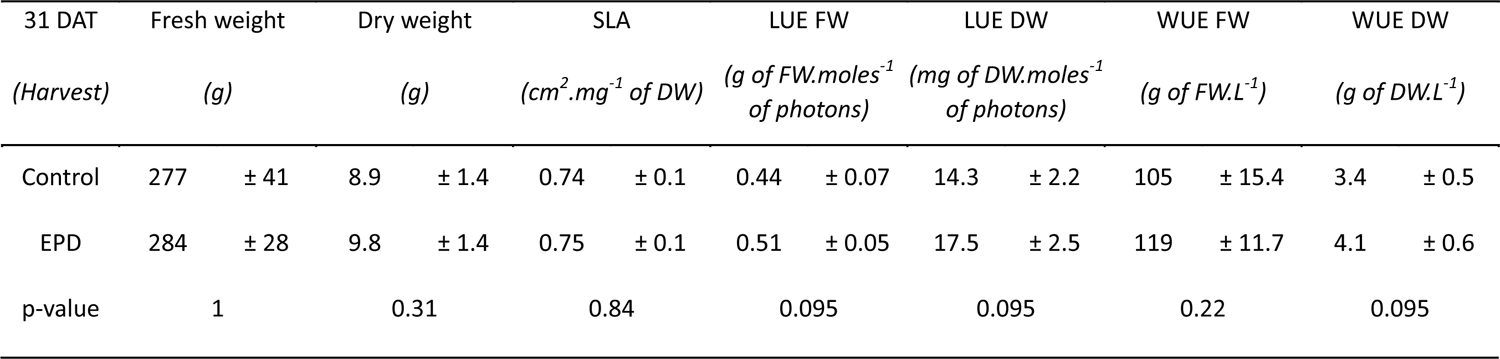
Yield and agronomical parameters measured in CT and EPD lettuces at 31 DAT (harvest). The ∑DLIs of CT lettuces were equal to 610 moles of photons.m^-2^, the ∑DLIs of EPD lettuces were equal to 584 moles of photons.m^-2^. P-values are a result of the non-parametrical pairwise Wilcoxon-Mann-Whitney test (n=5, α=0.05).

## Discussion

### The circadian rhythm of stomata regulates the variations of photosynthesis, whether it is tuned or exceptionally not tuned with the photoperiod

Throughout this experiment, net photosynthesis, stomatal conductance, PSII efficiency, maximum PSII efficiency of light-adapted lettuces and photochemical quenching showed daily variations. In both modalities, the net photosynthetic activity showed a sinusoidal pattern during the light period, consistent with the patterns already described in *Symbodinium sp.*, *G. polyedra*, *P. vulgaris*, *M. polymorpha* and *Z. aethiopica* (Hastings et al., 1961; Fredeen et al., 1991; Sorek et al., 2013; Cuitun-Coronado et al., 2022; Dias et al., 2022). However, the patterns between EPD and CT lettuces were different. In lettuces studied after a longer period of darkness, the mean net photosynthesis was 6.2% lower than the control. The loss of photosynthetic activity after an exceptionally longer period of darkness is consistent with the literature, as it has been showed that *A. thaliana* expression circadian rhythmicity of 24 hours (Col-0), 20.7 hours (*toc1-1* mutants) and 27.1-32.5 hours (*ztl-1* mutants) assimilate at least 40% less carbon when the photoperiod were not tuned with their circadian rhythm (Dodd et al., 2005).

In order to better describe the link between the photosynthetic activity and the circadian rhythm, we expressed the studied photosynthetic parameters according to the anticipation of the light period by the circadian rhythm (ZT-CR; figure 5), rather than according to the actual hours after the beginning of the light period (ZT; figure 2). Concerning the control, no changes occurred between ZT and ZT-CR-based representations, which validate that the photoperiod and the circadian rhythm were tuned (figure 2A&5A). However, for EPD lettuces, the conversion of ZT to ZT-CR induces a shift of 4 hours in photosynthetic parameters trends, as the darkness period was extended to 4 hours (figure 2B&5B). When the photosynthetic parameters are expressed based on ZT-CR, their variations throughout the light period are similar to the control (figure 2&5). It has to be noted that, whatever the representation, the stomatal conductance is always the first parameter to decrease. Stomatal conductance, *i.e.* stomatal opening, is already known to be under circadian control (Kreps and Kay, 1997). Moreover, we observed a synchronous decrease of net photosynthesis and stomatal conductance values. These data underline the link between photosynthesis and stomatal conductance which has already been described (Gorton et al., 1989). Multi-species meta-analysis approach has demonstrated a significant positive correlation between net photosynthesis and stomatal conductance (p-value < 0.001) (Gago et al., 2016). Indeed, stomatal conductance regulates the flow of incoming CO_2_ into the leaf, which may limit Rubisco carboxylase activity and therefore photosynthetic activity (Hennessey and Field, 1991). The correlation between net photosynthesis and stomatal conductance is also verified during the following light period, when the photoperiod and the circadian rhythm of lettuces are re-tuned (figure 3). Interestingly, the interrelation between the photosynthetic parameters within a light period remains unchanged whatever the modality, which suggests that the disruption of the circadian rhythm did not induce drastic changes in the physiology of the lettuces (Supplementary figure S1). Thus, variations in net photosynthetic activity depend on the circadian rhythm of stomatal conductance, whether the photoperiod and the circadian rhythm are tuned or exceptionally not tuned.

**Figure 5.**
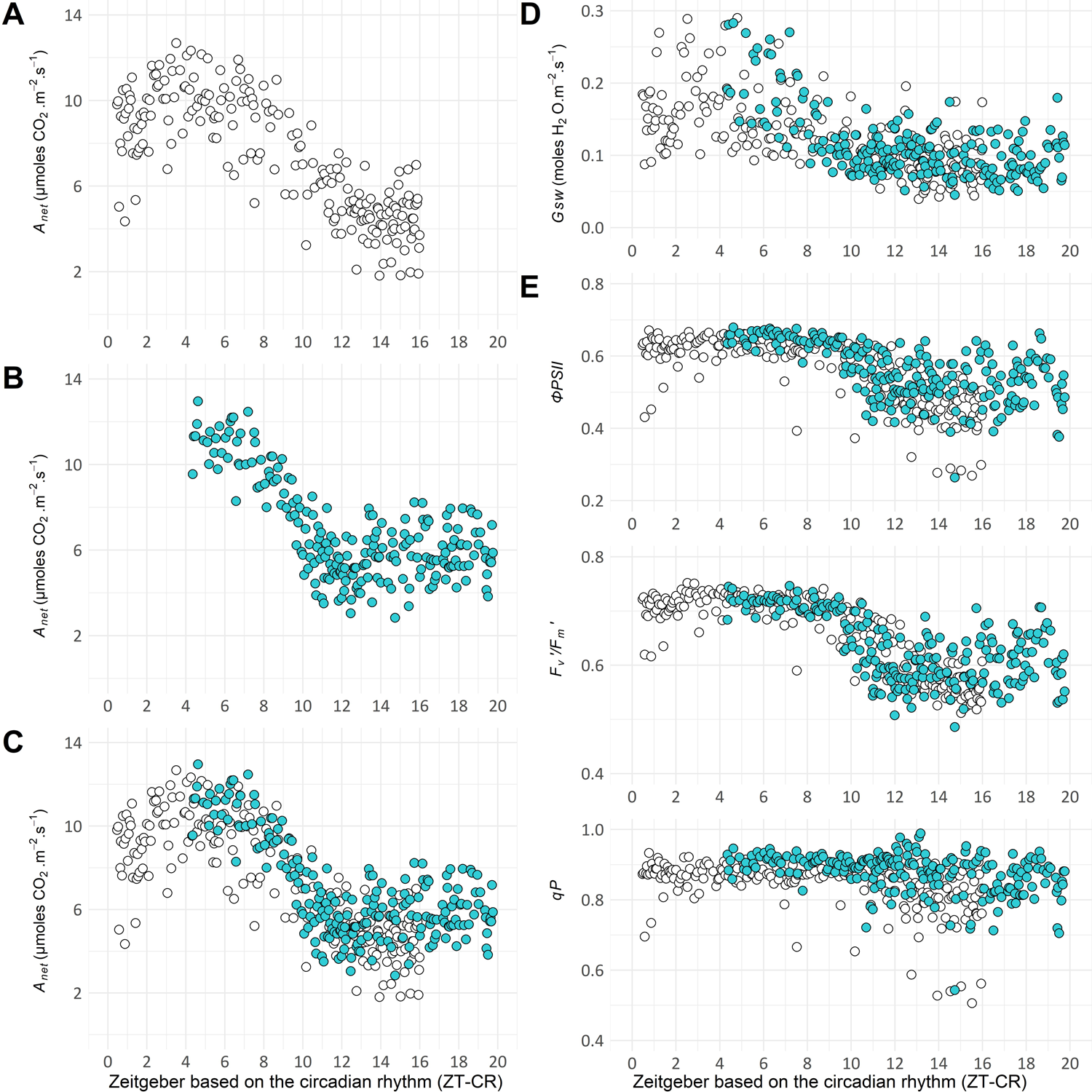
Photosynthesis (*A_net_*), stomatal conductance (*Gsw*), photosystem II efficiency (*ΦPSII*), maximum photosystem II efficiency of light-adapted plants (*F_v_’/F_m_’*) and photochemical quenching (*qP*) over 16 hours for CT (white circles) and EPD lettuces directly after the extend of the darkness period to 12 hours (cyan circles), and expressed as hours after the beginning of the light period based on the anticipation of the light period by the circadian rhythm (ZT-CR). (A) *A_net_* values for CT lettuces. (B) *A_net_* values for EPD lettuces after a 12 hours darkness period. (C) Comparison of *A_net_* values between CT and EPD lettuces. (D) Comparison of *Gsw* values between CT and EPD lettuces. (E) Comparison of *ΦPSII*, *F_v_’/F_m_’*, *qP* values between CT and EPD lettuces.

Light harvesting process indicators are decreasing later in the light period in comparison with the stomatal conductance. Photorespiration activity of the Rubisco, *i.e.* oxygenase activity, is known to play a key role in electron management (Kozaki and Takeba, 1996), particularly when its carboxylase activity is limited by the stomatal conductance. In our experiment, results suggest the stable efficiency of the photosystem II up to two hours after the closure of the stomata could be allowed by the photorespiration process. After two hours, the efficiency of the photosystem II decreased, which seems unlikely to be explained by the Rubisco activity, as it has been described as not subject to a circadian rhythmicity (Fredeen et al., 1991; Hollnagel et al., 2002). In our experiment, we noted a synchronous decrease of the different parameters related to the photosystem II efficiency (*ΦPSII*, *F_v_^’^/F_m_^’^*, *qP*), which is consistent with the fact that the decrease of the photosystem II efficiency may be linked to a decrease in the maximum capacity of the efficiency of the photosystem II, regulated by the circadian rhythm (Dodd et al., 2005; Dodd et al., 2014b). We can so hypothesize that this decrease is linked to the circadian rhythmicity of genes that are specifically regulating the efficiency of photosystem II. Indeed, Harmer et al. (2000) have studied the levels of genes related to photosystem II in *A. thaliana* and they observed a clear circadian rhythmicity of these genes under constant lighting conditions. More precisely, the transcription of genes encoding for chlorophyll binding proteins (LHCA and LHCB) and PSI and PSII reaction centers were regulated by the circadian rhythm and peaked at approximately ZT4 (Harmer et al., 2000). Further studies are therefore needed to better characterize the variations in photosystem II efficiency throughout the light period. Indeed, circadian variations in photosystem II efficiency are difficult to describe because of their dependence on the leaf spot and the species studied (Allen, 2000; Rascher et al., 2001).

### Photosynthesis is not impacted by the starch depletion at the end of the exceptionally long period of darkness

In lettuces, we have demonstrated that the photosynthesis is regulated by the circadian rhythm of stomatal conductance. In order to better characterize the consequences of this regulation, we assayed starch, sucrose, glucose and fructose contents. Starch content of lettuces after an exceptionally long period of darkness was 68% lower than the control. However, when the darkness period returned similar to the control, starch content of lettuces also returned similar to the control. After an exceptional darkness period of 12 hours, starch residual content of lettuces was equal to 12% of the initial stock, compared to 37% for the control after 8 hours of darkness. In *A. thaliana*, the residual starch content variates from 10 to 41% of the initial stock for darkness periods from 20 to 6 hours (Sulpice et al., 2014). In our experiment, the differences in starch residual contents cannot be explained by starch accumulation during the light period. Indeed, the light period was 16 hours for both modalities, which led to a similar starch accumulation, consistent with the literature as starch is accumulated linearly during the light period (figure 4D) (Gibon et al., 2004). However, the differences in residual starch contents can be explained by the adjustment of degradation rates during the darkness period. When the photoperiod and the circadian rhythm are tuned in *A. thaliana*, it has been measured that the starch degradation rate decreased from a maximum of 7.0 ± 0.9 µmoles C6_eq_.g^-1^ of FW.h^-1^ to 5.2 ± 0.3 µmoles C6_eq_.g^-1^ of FW.h^-1^ when the darkness period increased from 6 to 12 hours (Sulpice et al., 2014). The same trend, in which starch degradation rate decreases when the darkness period increases, has been observed as well in *A. thaliana* in the study of Gibon et al. (2004). In our experiment, when the photoperiod and the circadian rhythm are exceptionally not tuned, we measured equal starch degradation rates of 2.3 ± 1.0 mg. g^-1^ of DW.h^-1^ and 2.3 ± 1.1 mg. g^-1^ of DW.h^-1^ as well as 26 ± 11 mg. g^-1^ of FW.h^-1^ and 30 ± 11 mg. g^-1^ of FW.h^-1^ when the darkness period increased from 8 to 12 hours. Put together, these results suggest that the rate of starch degradation during a darkness period can only be adjusted when the photoperiod is tuned with the circadian rhythm. The link between circadian rhythm and starch has already been deeply characterized in *A. thaliana* (Gibon et al., 2004; Bläsing et al., 2005; Graf et al., 2010; Scialdone et al., 2013; Sulpice et al., 2014; Fernandez et al., 2017). In our study, we can suggest that this link between starch content and circadian rhythm is verified on lettuces and we measured that in lettuces, the mean starch degradation rate was approximately equal to 2.6 mg of starch. g^-1^ of DW.h^-1^ during the darkness period, which approximately corresponds to 27 µg of starch. g^-1^ of FW.h^-1^ during the darkness period.

Starch is degraded into sucrose, which might explain that the sucrose content pattern follows that of starch for control lettuces. This pattern was not repeated in both modalities, as sucrose content increases 48 hours after the lettuce undergoes a shift between the photoperiod and the circadian rhythm. Sucrose is known to be the main transportable form of carbon, which may induce variability in its content (Lemoine, 2000). Thus, the biosynthesis of sucrose, partitioning and transport need to be better characterized when the photoperiod and the circadian rhythm are not tuned. Glucose and fructose did not variate according to the photoperiod or according to the circadian rhythm. Moreover, no trehalose was found in our samples (data not shown). Therefore, these results suggest that starch is the best candidate as to be indicator of the circadian rhythm amongst the studied sugars.

In this experiment, we investigated the link between starch content and net photosynthesis. The depletion of starch after an exceptionally long period of darkness did not have any impact on net photosynthetic activity during the following light period. Today, we lack on information on this subject in the literature. However, the link between starch and maximum photosynthetic capacity has already been studied. Indeed, on *A. thaliana* and *O. sativa*, an increase in leaf starch content increases the photosynthetic capacity of the plant (Sun et al., 1999; Gibson et al., 2011). Starch is known to recycle inorganic pyrophosphate in C_3_ plants through the activity of the ADPglucose pyrophosphatase, which occurs in addition to the recycling by the photorespiration, and then plays a key role in the inhibition of photosynthesis related to the phosphate content (Ogren, 1984; Sharkey, 1985; Suzuki et al., 2021). Furthermore, a linear correlation between the amount of ^14^C incorporated into starch and net photosynthesis has been described, with a coefficient of correlation equal to 0.90 (Sun et al., 1999). However, starch content depends on starch precursors, such as glucose and fructose. In our experiment, a saturating light was selected for the control and allows no light-related limitations of the photosynthesis (Zhou et al., 2022), and thus of glucose and fructose biosynthesis. When lettuces were exposed to exceptionally long periods of darkness, the glucose and fructose contents were approximately 23% and 32% lower than the control respectively, probably link to a buffering of starch over-degradation compare to the control. Despite those differences in starch, glucose and fructose contents, we observed no inhibition of the net photosynthesis throughout the day, which is consistent with the fact that no direct feedback inhibition of the photosynthetic capacity is induced by photosynthesis end-products (Goldschmidt and Huber, 1992). Thus, it seems that starch content regulating the maximum photosynthetic capacity (Suzuki et al., 2021), but the starch-dependent limitation of net photosynthesis was not the main limiting factor in this study, likely less limiting than the stomatal conductance.

### The yield is not altered by a disruption of the circadian rhythm if the pool of carbohydrates is not exhausted

Fresh weight, dry weight, specific leaf area, water use efficiency and light use efficiency were similar for EPD lettuces and the control. However, the decrease of 6.2% in net photosynthesis during the light period following an exceptionally long period of darkness could have induced a reduction in the growth rate of lettuces. Previous studies have already showed that a photoperiod not tuned with the circadian rhythm induces a reduction of the photosynthetic activity. Indeed, Dodd et al. (2005) showed that *A. thaliana* expressing a circadian rhythmicity of 24 hours (12/12) fixed 42% and 47% more CO_2_ with a day of 24 hours than with days of 28 hours (14/14) or 20 hours (10/10) respectively. The same pattern has been observed when *A. thaliana* expressing a circadian rhythm of 20 hours or 28 hours were placed in longer (28 hours) or shorter (20 hours) photoperiods respectively (Dodd et al., 2005). Moreover, in *A. thaliana*, Graf et al. (2010) have measured a decrease of the fresh weight of 37% when the length of the day length reaches 28 hours (14/14) instead of 24 hours (12/12). In the literature, the decrease in photosynthetic activity is higher than in our experiment, consistent with the proportion of the disruption of the circadian rhythm.

In our study, the net photosynthesis has been assumed to be solely limited by the circadian rhythm of the stomatal conductance. However, in previous studies, long periods of darkness might have induced a transitory depletion state of carbohydrates at the end of the darkness period, which has been assumed to inhibit growth (Gibon et al., 2004; Graf et al., 2010). Thus, we can propose that the yield of lettuces was not altered by the disruption of the circadian rhythm because the pool of carbohydrates was not exhausted at the end of the darkness period, notably glucose and fructose. To summarize, we can propose that the threshold for growth-limiting glucose and fructose content is less than 75 mg. g^-1^ of DW and 100 mg. g^-1^ of DW respectively in our experimental conditions. Finally, we demonstrated that the photosynthetic activity and the carbohydrates management are not limited by the same factors, suggesting that growth regulation must take both into account when photoperiod is exceptionally not tuned with the circadian rhythm. Interestingly, the exceptional disruptions of the circadian rhythm allowed to save 5% of lighting time compared to the control, which justifies that circadian rhythm has to be considered to reduce the energy consumption of indoor farming facilities while producing a marketable yield.

## Conclusion

In conclusion, this work demonstrates that the circadian regulation of the stomatal conductance is the main limiting factor of net photosynthesis when the photoperiod is exceptionally not tuned with the circadian rhythm. In the conditions of this study, the photosynthetic activity during the light period following and exceptionally long period of darkness decreased of 6.2% compared to the control. Consequently, the management of carbohydrates was impacted and glucose and fructose contents were significantly lower for lettuces under an exceptionally long period of darkness. Starch and sucrose contents were significantly lower than the control after a longer period of darkness but returned similar to the control at the end of the following photoperiod. Starch had been previously assumed to be involved in the maximum photosynthetic capacity. However, the differences in starch contents according to the darkness periods did not inhibit the net photosynthesis of the following light period. Moreover, no differences in yield, morphology or efficiency were found between both modalities. Therefore, we assumed that growth cannot be directly impacted by the exceptional disruptions of the circadian rhythm through net photosynthesis. This inhibition occurs through carbohydrates, and the thresholds at which growth will be altered need be further characterized. In this work, disrupting the circadian rhythm saved 5% of lighting time compared with the control, which can be converted into energy savings. To conclude, this mutant-free experiment supports the idea that the circadian rhythm needs to be considered when the photoperiod is modified, which is a fast-growing topic for reducing energy consumption in indoor farming. Thus, further studies are required to better characterize the photosynthesis-carbohydrates-growth continuum under conditions where the photoperiod is tuned, not tuned or exceptionally not tuned with the circadian rhythm.

## Acknowledgments

We thank Doriane Dumont for sharing her experiences and advices on starch assays. We also thank Louis Ramade for his help during the experiments.

